# Pre-Assembly NGS Correction of ONT Reads Achieves HiFi-Level Assembly Quality

**DOI:** 10.1101/2024.07.12.603260

**Authors:** Evgeniy Mozheiko, Heng Yi, Anzhi Lu, Heitung Kong, Yong Hou, Yan Zhou, Hui Gao

## Abstract

**Background:** Recently developed hybrid assemblies can achieve Telomere-to-Telomere (T2T) completeness of some chromosomes. However, such approaches involve sequencing a large volume of both Pacific Biosciences high-fidelity (HiFi) and Oxford Nanopore Technologies (ONT) sequencing reads. Along with this, third-generation sequencing techniques are rapidly advancing, increasing the available length and accuracy. To reduce the final cost of genome assembly, here we investigated the possibility of assembly from low-coverage samples and with only ONT corrected by Next-Generation Sequencing (NGS) sequencing reads. We demonstrated that ONT-based assembly approaches corrected by NGS can achieve performance metrics comparable to more expensive hybrid approaches based on HiFi sequencing.

**Results:** We investigated the assembly of different chromosomes of HG002 and the low-coverage performance of state-of-the-art hybrid assembly tools, including Verkko and Hifiasm, and ONT-based assemblers Shasta and Flye. Our study shows that even with the best NG50, Hifiasm has many misassemblies in the centromere region, raising doubts about its T2T capabilities. Additionally, ONT-based assemblies perform well at low coverage, offering a cost-effective solution. To achieve better assembly quality, we evaluated the performance of different NGS technologies across various aspects of hybrid genome assembly, including pre-assembly correction, haplotype phasing, and polishing, and found them to be similarly effective. For the NGS technology stLFR, we proposed two-round assembly methods that utilize stLFR linked-read data to achieve assembly phasing performance comparable to that obtained with trio data.

**Conclusion:** We found that assemblies based on ONT corrected by NGS can match or exceed HiFi-based assemblies in accuracy. It appears that no assembly tool investigated here can achieve T2T completeness, even with 50X coverage. Therefore, we recommend using a combination of regular R9 or R10 simplex ONT reads and accurate NGS reads for assembly without aiming for T2T completeness. These ONT-based assemblies perform well even at low coverage, offering a cost-effective solution.

## Background

Genome assembly, the process of reconstructing the complete genomic sequences from fragmented sub-sequences, is a critical step in genomics research. Recent advancements in sequencing technologies have seen the rise of third-generation sequencing data such as ONT and HiFi [1–3], providing long-read sequencing data that has dramatically improved the quality of genome assemblies [4–16]. The rapid evolution of third-generation sequencing techniques has paved the way for hybrid assembly algorithms [4–6]. These sophisticated methodologies combine the accuracy of short reads from NGS, such as MGI or Illumina [17–19], with the structural information provided by long-read sequencing data to generate high-quality and complete genome assemblies. Despite their effectiveness, these hybrid approaches typically involve sequencing a large volume of both long accurate HiFi and ultra-long ONT reads, which can be resource-intensive [20].

Currently, most hybrid assembly algorithms rely heavily on HiFi reads [4–6]. Error-prone ONT reads are primarily used for filling gaps and aren’t involved in the assembly consensus where there is enough HiFi coverage [4–6]. Moreover, before-assembly self-correction is carried out without involving the ONT, which doesn’t fully utilize the potential of ultra-long reads [4–6]. The reason why ONT are not used in self-correction explained as they are too error prone for this purpose. The recent achievement in ONT self-correction opened this possibility, but only with using machine learning algorithms [21] or without achievement HiFi level of assembly quality [22]. The other way to reduce error rate of ONT is correction by short accurate NGS reads. The tools like Ratatosk [23] are used for ONT correction with NGS reads, which allow to achieve better accuracy, but still less than HiFi or NGS [23].

Because of ONT accuracy can not achieve HiFi accuracy even after NGS correction, therefore corrected ONT reads cannot be used in HiFi-based tools instead of HiFi. Even with absence of availability of using ONT reads corrected by NGS within HiFi-based tools, there is still the possibility to use them in ONT-based tools. The performance of corrected of ONT by NGS are not wide evaluated with State-of-the-art ONT-based assembly tools like Shasta or Flye. In this study we close this gap in literature and demonstrated that ONT-based assembly tools can achieve same assembly accuracy when using ONT reads corrected by NGS as input.

Hybrid assembly tools find NGS and Hi-C applications for genome phasing [24]. This makes it possible to obtain an almost complete assembly of diploid genomes. Nevertheless, this often requires large input coverages, more than 50X of sequencing reads and the parameters of some of genome assembly tools such as Verkko are not adapted to the use of low coverage [4]. One of the ways to reduce the final cost of genome assembly is to use lower coverage. At the same time, most assembly tools are focused on optimizing run time or improving the performance of various quality metrics with a high coverage of more than 50X [4–6, 8, 10, 16]. In its turn, assemblies with coverages lower than 30X are poorly investigated. Another approach to cost reduction involves exclusively utilizing ONT and NGS while foregoing the more costly PacBio HiFi data. The primary challenge to address is the acquisition of accurate corrected reads, where self-correction or hybrid correction may serve as viable solutions. Additionally, substituting NGS data from the pricier Illumina platform with data from a more cost-effective yet equally precise NGS platform, such as MGI, could be considered. Furthermore, during the haplotyping of assembly results, trio NGS data could be replaced with linked-read data. In this article, we will explore each of these aspects and evaluate the performance of these substitutions, which could facilitate broader adoption of high-quality assembly work.

The difference between the main NGS technologies MGI and Illumina in the context of hybrid assembly tools also remains limited [4– 6, 25], which does not allow choosing the most optimal choice of such data for research. Here we carried out comparative analysis of these technologies’ performance and their potential for effectively reducing the final cost of genome assembly. Moreover, we estimated the possibility of obtaining contiguous assemblies with only NGS + ONT sequencing reads. The main goal of this study is to show that the development of ultra-long read technology could help to resolve complex genomic regions and at the same time substantially reduce the assembly cost due to low coverage. Quality evaluation metrics such as NG50, QV score, Hamming and Switch error were utilized to benchmark the assemblies [26–28].

## Methods

### Genome sequencing

The library preparation was performed with an automated liquid handling platform, MGISP-960 (MGI), using MGIEasy FS PCR-Free DNA Library Prep Kit V1.2 (MGI) according to the manufacturer’s protocol. Briefly, the samples were fragmented at 30□°C for 15□min and 65□°C for 15□min. The fragmentated DNAs were size selected in between 450 to 600bp. end-repaired and A-taling were performed at 14°C for 15 min, at 37°C for 25 min and at 65°C and 15 min. The end-repaired DNAs were indexed at 25°C for 10 mins with adapter-barcode oligos (Number 89 to 102) in the kit. The adapter-ligated DNAs were cleaned up with MGIEasy DNA Clean beads Kit (MGI) and denatured at 95°C for 3 mins to obtain single-strand DNAs. The Circularization Enzyme Mix in MGIEasy FS PCR-Free DNA Library Prep Kit V1.2 was used to form the circularized-DNAs at 37°C for 10 mins. DNB (DNA Nano Ball) generation was performed with MGISP-960 (MGI) using DNBSEQ-T10×4 RS High-throughput Sequencing Kit according to the manufacturer’s protocol at 30°C for 25 mins. Finally, the DNBs were sequenced on the DNBSEQTM T10 sequencing platform (MGI) with 150 bp read1 and 150 bp read2.

### WGS data analysis

Human samples from Ashkenazim Trio (HG002, HG003, HG004) were aligned to the CHM13 reference, detailed coverage information was showed in supplementary tables. CHM13 FASTA was indexed with Minimap and BWA with parameters ‘minimap2 - ax map-pb -t 32’ for HiFi, ‘minimap2 -x map-ont -t 32’ for ONT and default parameters for BWA index. ONT and HiFi were aligned with Minimap2 with corresponding indexes, and NGS with BWA with parameters ‘BWA mem -t 32’ [29, 30]. After this all reads aligned to chromosomes from 1 to 22, were sorted, indexed and downsampled with SAMtools ‘samtools sort -@ 32, samtools view -@ 32 -b, samtools index -@ 32, samtools fastq’ [30]. SeqKit was used to remove duplicates from reads downsampled to distinct chromosome ‘seqkit rmdup -s’ [31]. Ready FASTQ files with only reads from distinct chromosome were subsampled with Seqtk into different coverages from 10X, 20X, 30X and 50X ‘seqtk sample -s100’ [31].

### Assembly tools

Here we tested assembly tools Verkko, Hifiasm, LJA, Shasta and Flye. All these tools were tested with different chromosomes and with different coverages: 10X, 20X, 30X, 50X. Hybrid assembly tools Verkko and Hifiasm were launched in haploid mode with default parameters. LJA was launched with default parameters. For the Ratatosk + Shasta and Ratatosk + Flye assemblies, we first corrected the ONT reads using NGS data with Ratatosk, employing the command Ratatosk correct -v -c 32. We then converted non-{A, C, G, T} characters using the Ratatosk Python script replaceIUPAC.py. All assembly tool commands used in this study are listed in Supplementary Table 6. We utilized PEPPER for ONT-polishing and Pilon for NGS-polishing of the assemblies, both with default parameters. To calculate the empirical accuracy of ONT reads corrected by Ratatosk, we first aligned the corrected ONT reads using minimap2 and then calculated the empirical accuracy of each read using the stats_from_bam tool from Pomoxis with default parameters https://github.com/nanoporetech/pomoxis.To assess assembly phasing with different NGS technologies we compared single-tube long fragment read (stLFR) [32], and MGI and Illumina trio. The main ideas of ONT + stLFR assembly are using phasing information from stLFR and two assembly rounds to make distinct assembly for every haplotype. The first round is a Shasta assembly with default parameters and BLR [33] with Shasta assembly and stLFR reads as an input with default parameters. The result of the first round is a phased Variant Call Format (VCF) file based of de novo Shasta assembly and stLFR reads. The second round is a haplotype binning of ONT and stLFR reads with this phased VCF, correction of ONT reads with stLFR separately for each haplotype with Ratatosk with default parameters, and haploid assembly with Flye [13] with default parameters.

For benchmarking assemblies from different tools, we calculated NG50, Contig Count, k-mers based completeness and QV score. NG50 and Contig Count were received from QUAST statistics [26]. QUAST was run with reference mode with CHM13 reference and other defaults parameters. K-mers based completeness and QV score were received from Merqury [27] with defaults parameters.

Phasing statistics such as Switch and Hamming error were calculated in MGI and Illumina as trio data comparison, and for this purpose we used the ‘yak trio eval’ command with default parameters.

Meryl databases for QV evaluation in MGI and Illumina as Meryl databases comparison were calculated with command ‘meryl k=31 threads=32 count’ for every type of sequencing reads. After this, received databases were combined with command ‘meryl union-sum’.

## Results

Modern assembly tools utilize hybrid approaches that incorporate the use of both long, accurate HiFi reads and ultra-long (UL), error-prone ONT reads simultaneously in a single assembly to achieve near telomere-to-telomere completeness. We suggest that error correction of ONT or ONT UL (>100 kb) reads with R9 chemistry by NGS short reads before assembly can lead to similar results. To get a more comprehensive view, we assembled each chromosome of HG002 separately with hybrid HiFi-based and corrected ONT-based assembly tools. Additionally, we evaluated short NGS read technologies MGI, Illumina and stLFR in the NGS + ONT assembly.

### ONT based and hybrid assemblers

We tested different ONT-based and HiFi-based assembly tools on human samples from the Ashkenazim Trio [34] (Supplementary Table 1). We used standard size and Ultra Long (UL) > 100kb ONT reads with R9 chemistry. Somatic chromosomes from HG002 were assembled with hybrid HiFi-based assembly tools Verkko, Hifiasm and LJA, and ONT-based assembly tools Shasta and Flye. It is knowns that ONT reads do not have the same high accuracy as HiFi or NGS, and only the latest R10 ONT libraries can reach Q30 [25, 35]. One of the ways to improve the accuracy of an ONT is to correct errors before assembly using correction tools such as Ratatosk which allow to polish ONT with NGS reads [23]. This helps to remove most of errors, reach 99% accuracy and at the same time use only NGS and ONT in hybrid assembly. Correction before assembly results in an improvement of all tested assembly quality statistics (Supplementary Figure 1). The most substantial improvements are in QV, completeness, and indels per 100kb quality metrics. Here we assessed the possibility of ONT corrected by NGS to be assembled with a similar performance as hybrid HiFi + ONT approaches (Figure 1, Supplementary Figure 2).

**Figure 1.**
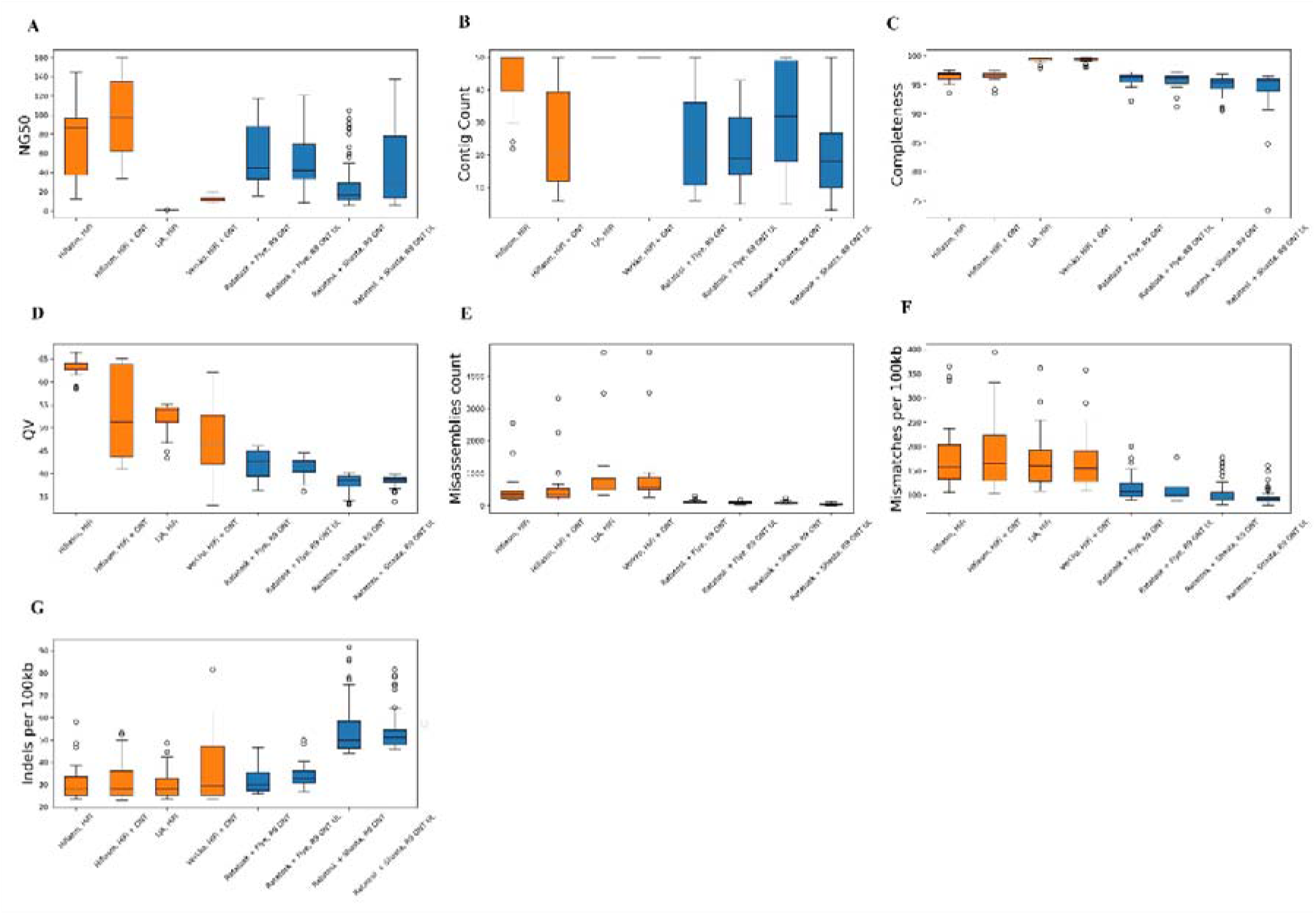
Quality statistics distribution of assemblies on chromosomes 1 to 22 of HG002. Blue boxplots for ONT + NGS based assemblies, orange for HiFi or HiFi + ONT based assemblies. **A**. NG50, **B**. Contig Count, **C**. K-mer based completeness, **D**. QV score. **E**. Overall count of misassembles **F**. Mismatches per 100kb **G**. Indels per 100kb.

Hifiasm has the best NG50 and QV here (Figure 1 A, D), but other HiFi-based tools, LJA and Verkko, demonstrated lower NG50 than corrected ONT-based assembly approaches (Figure 1 A). Despite higher NG50 and QV, Hifiasm and other HiFi-based tools showed up to 10 times more misassemblies (Figure 1 E) and worse mismatch statistics (Figure 1 F). In turn, HiFi-based methods performed better with indel statistics than Ratatosk + Shasta. Ratatosk + Flye with R9 ONT have similar indel statistics as HiFi-based methods and slightly worse with R9 ONT UL (Figure 1 G). Between the two corrected ONT-based assembly tools, Ratatosk + Shasta and Ratatosk + Flye, Ratatosk + Flye demonstrated better performance across all statistics except for misassemblies and mismatches. In summary, ONT-based tools are worse with NG50 and QV, but better with contig count, misassemblies and mismatches, and have similar or slightly worse k-mer-based completeness and indel statistics than corrected HiFi-based methods.

ONT UL is more expensive than regular ONT and is usually used only to assemble up to T2T completeness in combination with other sequencing platforms such as HiFi [4, 36]. Nevertheless, no assembly tool in this study can achieve T2T completeness even with 50X coverage (Supplementary Figures 3-9). This is due to the presence of numerous misassemblies within the centromere region, even when assembling the centromere with one contig. We investigated the assembly of chromosomes 7, 9, 10, 11, 12, 16, and 20, where Hifiasm was substantial better with NG50 (Supplementary Figure 2) and found that Ratatosk + Flye with R9 ONT demonstrated one-contig contiguity outside of centromeric regions and partially assembled centromeric regions (Supplementary Figures 3-9). Therefore, Hifiasm’s advantage in NG50 remains under discussion.

Generally, genome assembly tools recommend a coverage of 30X or preferably 50X [4–16]. However, these studies did not publish their performance at lower coverages (Supplementary Table 2). Therefore, to address this in this study input sequencing data was adjusted to simulate different mean genome coverages of 10X, 20X, 30X and 50X. For each coverage we assembled each chromosome separately and plot average values of assembly quality metrics for every tool (Figure 2). Corrected ONT-based approaches occurred more stable at low coverages than HiFi based tools (Figure 2). This results in better NG50 and indels statistics of Ratatosk + Flye at 10X coverage than all assembly methods observed in this study (Figure 2 A, G). It is important to note that even with 10X coverage Flye demonstrates a good level of NG50 across all chromosomes (mean value over 40Mb), and completeness over 95% (Supplementary Figure 10). Therefore, low coverage assembly performance is a second advantage of corrected ONT-based approaches over HiFi-based methods.

**Figure 2.**
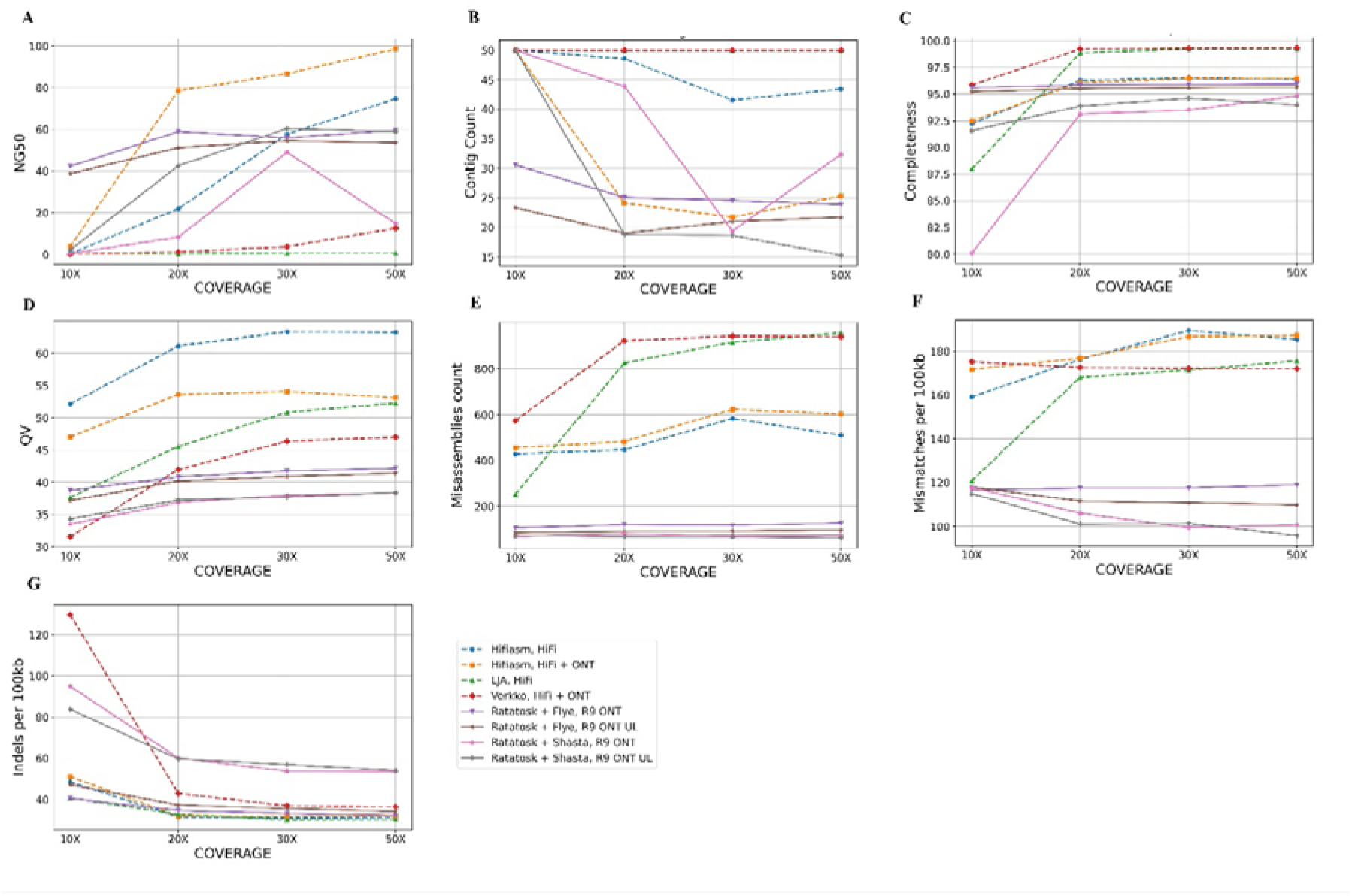
Quality statistics of assembly tools with low coverage. Each chromosome of HG002 was assembled and plotted mean values of assembly quality statistics. Solid lines are for ONT + NGS based assemblies, dashed is for HiFi or HiFi + ONT based assemblies. **A**. NG50, **B**. Contig Count, **C**. K-mer based completeness, **D**. QV score. **E**. Overall count of misassembles **F**. Mismatches per 100kb **G**. Indels per 100kb.

### MGI and Illumina in hybrid assembly

We investigated the possibility of using Trio NGS + ONT or NGS + ONT as an alternative to Trio NGS + HiFi + ONT or HiFi + ONT respectively for hybrid assembly. The two most used NGS technologies MGI and Illumina have not been distinctly compared in this context. Therefore, to determine which NGS technology has better performance in a hybrid NGS + ONT assembly, we compared MGI and Illumina as trio data for haplotype phasing, in correction before-assembly, in polishing after assembly and as Meryl databases for QV score evaluation. We tested two samples for parental data (HG003, HG004) and two samples for HG002 for MGI and the same for Illumina.

#### NGS as trio in haplotype phasing

Hybrid assembly tools like Verkko and Hifiasm utilize NGS as trio data for haplotype phasing [4, 5]. Among these two genome assembly tools, Hifiasm showed better haplotype phasing performance [4]. Therefore, we chose Hifiasm for testing MGI and Illumina reads as trio input. Two MGI and two Illumina samples from the Ashkenazim trio were tested, and MGI demonstrated similar or slightly better quality control statistics (Table 1, Supplementary Table 3). Also, for a more comprehensive view, we compared Illumina with another MGI NGS technology, stLFR. stLFR is a technology simulating long reads, adding the barcode information to each subfragment of the long DNA molecules which could achieve 100kb [37]. The advantage of this approach is that it combines the benefits of both long-read and short-read approaches while maintaining the accuracy of short reads and store the long-range information. stLFR does not require sequencing of parental genomes and shows similar phasing statistics to trio-based assembly methods (Table 1).

**Table 1.**
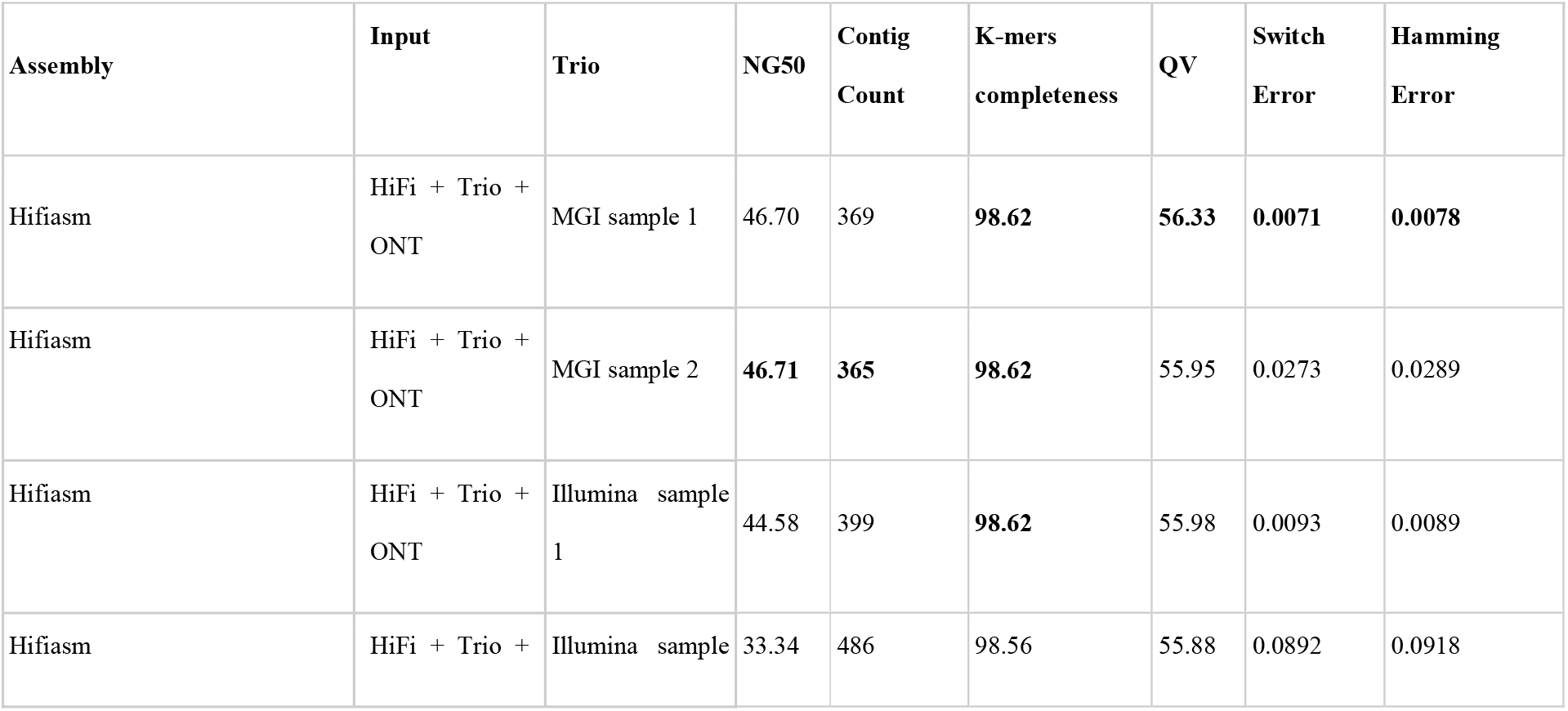
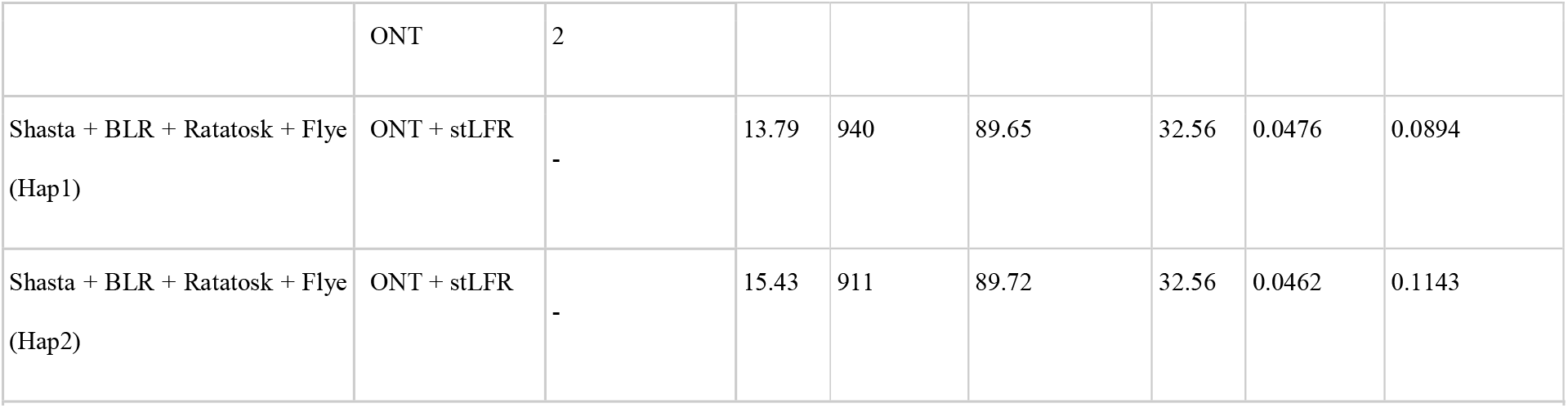
Quality statistics of Hifiasm with MGI or Illumina as a Trio. HG002-HG004 trio assembled with Hifiasm in trio mode - HiFi + ONT + Trio (MGI / Illumina). The whole genome was utilized for this assemblies.

#### NGS in error correction prior assembly

Another application of NGS in hybrid assembly is the correction of error-prone ultra long ONT reads. Before-assembly and after-assembly correction are the two most common approaches [16, 23, 38, 39]. In before assembly case, we selected Ratatosk, one of the recent pre-assembly correction tools, to compare the MGI and Illumina NGS technologies [23]. Ratatosk is based on a compacted and colored de Bruijn graph built from accurate short NGS reads. Here we performed correction of ONT reads with two MGI and two Illumina samples (HG002). After correction, we empirically estimated the accuracy of corrected ONT reads. Correcting ONT reads using either MGI or Illumina resulted in a similar distribution of accuracy using either MGI or Illumina resulted in a similar median accuracy: 98.66, 98.46 for MGI and 98.49, 98.57 for Illumina samples (Supplementary Table 4, Supplementary Figure 11).

#### NGS in error correction after assembly

To compare MGI and Illumina in the context of after-assembly correction, we choose PEPPER + Pilon polishing approach. This is a hybrid polishing approach recommended in the Shasta article [16, 40]. PEPPER + Pilon starts from ONT polishing and ends with short reads NGS polishing. Therefore, we compared MGI and Illumina performance in the context of after-assembly PEPPER + Pilon polishing (Table 2). In this case, MGI and Illumina also demonstrated similar results.

**Table 2.** QV of polished Shasta assembly. HG002 ONT reads assembled with Shasta and polished with PEPPER + Pilon. Only chromosome 20 was utilized for this assemblies and polishing.

| NGS | Sample | Polishing | QV |
| --- | --- | --- | --- |
| - | - | - | 26.92 |
| - | - | PEPPER | 29.19 |
| Illumina | 1 | PEPPER + Pilon | 31.71 |
| Illumina | 2 | PEPPER + Pilon | 31.72 |
| MGI | 1 | PEPPER + Pilon | 31.69 |
| MGI | 2 | PEPPER + Pilon | 31.46 |

#### NGS as Meryl databases in QV evaluation

Quality evaluation is an important step in genome assembly. One of the metrics used for this purpose is the QV score and k-mers based completeness [27]. However, our results suggest that relying solely on sequences from the same sample for QV evaluation may not provide a comprehensive representation and can yield varying results depending on the input data (Table 3), it is proved that QV estimates based on different input reads are not strictly comparable. To further investigate this, we constructed Meryl [27] databases for QV evaluation using combinations of reads from different sequencing technologies. QV evaluation of Hifiasm assembly with HiFi, ONT or NGS Meryl databases showed up to 2 times differences. Specifically, the use of HiFi as a k-mer Meryl database resulted in a QV score approximately twice as high as that achieved with ONT. In contrast, MGI and Illumina showed close results to each other in this context (Table 3). Moreover, the combination of the different sequencing technologies only led to higher QV scores. Therefore, in the context of QV evaluation MGI and Illumina demonstrated similar performance.

**Table 3.** Different Meryl databases in QV evaluation and k-mers based completeness of Hifiasm assembly. HG002-HG004 trio assembled with Hifiasm in trio mode - HiFi + ONT + Trio MGI. QV evaluation with Merqury. Only chromosome 20 was utilized for this assemblies.

| Tools | Data base for QV | Base-level error rate:<br>QV | K-mer-based Completeness (%) |
| --- | --- | --- | --- |
| Hifiasm | HiFi | 58.85 | 99.65 |
| Hifiasm | ONT | 28.07 | 65.83 |
| Hifiasm | MGI / Illumina | 33.02 / 33.93 | 99.30 / 98.95 |
| Hifiasm | MGI / Illumina + HiFi | 58.96 / 59.03 | 99.62 / 99.49 |
| Hifiasm | MGI / Illumina + ONT | 35.65 / 37.08 | 97.84 / 97.64 |
| Hifiasm | MGI / Illumina + HiFi + ONT | 59.65 / 59.73 | 98.82 / 98.84 |

## Discussion

The use of a genome assembly approach based solely on NGS and ONT sequencing data may lead to a reduction in assembly costs. This study demonstrated that current ONT-based assembly tools can achieve performance similar to hybrid HiFi + R9 ONT UL genome assembly methods when ONT data is corrected by NGS prior to assembly. R10 ONT was not tested here, but we assume that all findings in this work can be applied to this more recent and accurate R10 ONT technology. Additionally, we recognize that the chromosome extraction process may introduce biases in the separate genome assemblies of extracted chromosomes. However, given that genome assemblers often use less stringent parameters for mapping, we suggest that chromosome extraction will not have a significant impact on the assembly of distinct chromosomes.

Although ONT-based approaches showed worse assembly NG50 and QV metrics, we found that HiFi-based methods, such as Hifiasm, exhibited significantly more misassemblies and mismatches. Consequently, the cost of merging contigs with Hifiasm is associated with a higher number of assembly errors. This low assembly quality within centromere regions leading us to question the T2T assembly capabilities of Verkko and Hifiasm with 50X coverage. Based on this study results, without this advantage in NG50, HiFi-based assembly tools Hifiasm, Verkko and LJA could be replaced by cheaper methods based on simplex ONT and NGS.

Corrected ONT-based approaches, such as Ratatosk + Flye, proved to be more stable at low coverages (less than 30X) and outperform all other tools investigated in this study at a coverage of 10X. Additionally, the NG50 of Ratatosk + Flye did not experience a sharp drop at 10X coverage as other tools and maintained a good level of other assembly quality metrics. In contrast, Ratatosk + Shasta also showed more stable results than HiFi-based methods but worse than Ratatosk + Flye. Given that Ratatosk + Flye with R9 ONT showed similar performance to R9 ONT UL, we suppose that Ratatosk + Flye with R9 ONT and NGS is the best choice for low coverage assembly.

We found that the hybrid HiFi + R9 ONT UL assembler Verkko and the HiFi-based LJA could not achieve even 20Mb NG50 (Figure 1A, Figure 2A). This aligns with the original Verkko article, where Verkko’s contiguity is several times worse without parental data [4]. Therefore, corrected ONT-based approaches outperform Verkko and LJA in all assembly quality statistics investigated here, except for completeness and QV. The high completeness and QV of Verkko and LJA are likely due to their k-mer-based assembly approach, specifically the Multiplex de Bruijn Graph [4, 41].

In this study, we examined the performance of the three NGS technologies: MGI, stLFR and Illumina, across four distinct aspects of genome assembly: as trio data in haplotype phasing, in the correction of ONT reads, assembly polishing, and as Meryl databases for QV evaluation. In all these aspects, MGI and Illumina demonstrated similar performance. ONT + stLFR also showed similar haplotype phasing statistics as a trio MGI and Illumina NGS samples. Therefore, stLFR could be considered as an alternative to the Trio data for the phased assembly. The second advantage of stLFR is that it could be used for before-assembly error correction of ONT reads and provide phasing information at the same time. Additionally, we created a Snakemake workflow that supported this research. This workflow contains the best practices of genome assembly and assembly evaluation and is available on GitHub https://github.com/MGI-EU/assembly_workflow.

## Conclusions

In this study, we evaluated low-coverage ONT sequences corrected by NGS as input for ONT-based assemblers and compared the results with those from HiFi-based assembly methods. Tools like Verkko and Hifiasm claim to achieve T2T assembly using both HiFi and UL ONT reads, but they measure success by having a single contig per chromosome. Our study shows that HiFi-based Hifiasm has the best NG50 here. However, even in the case of assembling a distinct chromosome with a single contig, Hifiasm has many misassemblies in the centromere region, raising doubts about its T2T capabilities. We found that ONT assemblies corrected by NGS can match or exceed HiFi-based methods in accuracy. Therefore, we recommend using a combination of regular R9 or R10 simplex ONT reads and accurate NGS reads for assembly without aiming for T2T completeness. Additionally, ONT-based assemblies perform well even at low coverage, offering a cost-effective solution.

## List of abbreviations

T2T: Telomere-to-Telomere
HiFi: Pacific Biosciences high-fidelity
ONT: Oxford Nanopore Technologies
NGS: Next-Generation Sequencing st
LFR: single-tube long fragment read
VCF: Variant Call Format
UL: ultra-long > 100kb

## Declarations

### Ethics approval and consent to participate

Not applicable.

### Consent for publication

Not applicable.

## Availability of data and materials

Data used in this study were obtained from publicly available sources (supplementary Table 5) and new generated data also available in the CNGB Nucleotide Sequence Archive (CNSA: https://db.cngb.org/cnsa; accession number CNP0004858). Snakemake workflow for assembly and assembly quality control is available at https://github.com/MGI-EU/assembly_workflow.

## Author contributions

Y.Z. conceived the problem and designed all detailed studies. H.G., A.Z.L and H.T.K performed the library preparation and sequencing. E.M. and H.Y. performed bioinformatics analysis. Y.Z., E.M., Y.H., H.G. and H.Y. coordinated the resources and facilitated insightful discussions. Y.Z., H.G., and Y.H. supervised the work. Y.Z., E.M. and H.Y. wrote the manuscript. E.M and H.Y. developed the analysis pipeline and published it on the GitHub. All authors read and approved the manuscript for submission.

## Acknowledgments

We sincerely thank the technical support provided by China National GeneBank.

## Supplementary Information

Supplementary Tables.xlsx – supplementary tables, data sources. Supplementary Figures.pdf – supplementary figures.

## References

[1] Logsdon GA, Vollger MR, Eichler EE. Long-read human genome sequencing and its applications. Nat Rev Genet 2020; 21: 597–614.

[2] Wenger AM, Peluso P, Rowell WJ, et al. Accurate circular consensus long-read sequencing improves variant detection and assembly of a human genome. Nat Biotechnol 2019; 37: 1155–1162.

[3] Jain M, Koren S, Miga KH, et al. Nanopore sequencing and assembly of a human genome with ultra-long reads. Nat Biotechnol 2018; 36: 338–345.

[4] Rautiainen M, Nurk S, Walenz BP, et al. Telomere-to-telomere assembly of diploid chromosomes with Verkko. Nat Biotechnol. Epub ahead of print 16 February 2023. DOI: 10.1038/s41587-023-01662-6.

[5] Cheng H, Jarvis ED, Fedrigo O, et al. Haplotype-resolved assembly of diploid genomes without parental data. Nat Biotechnol 2022; 40: 1332–1335.

[6] Jarvis ED, Formenti G, Rhie A, et al. Semi-automated assembly of high-quality diploid human reference genomes. Nature 2022; 611: 519–531.

[7] Nurk S, Walenz BP, Rhie A, et al. HiCanu: accurate assembly of segmental duplications, satellites, and allelic variants from high-fidelity long reads. Genome Res 2020; 30: 1291–1305.

[8] Ruan J, Li H. Fast and accurate long-read assembly with wtdbg2. Nat Methods 2020; 17: 155–158.

[9] Bankevich A, Bzikadze A V., Kolmogorov M, et al. Multiplex de Bruijn graphs enable genome assembly from long, high-fidelity reads. Nat Biotechnol 2022; 40: 1075–1081.

[10] Chin C-S, Peluso P, Sedlazeck FJ, et al. Phased diploid genome assembly with single-molecule real-time sequencing. Nat Methods 2016; 13: 1050–1054.

[11] Vaser R, Šikić M. Time- and memory-efficient genome assembly with Raven. Nat Comput Sci 2021; 1: 332–336.

[12] Koren S, Walenz BP, Berlin K, et al. Canu: scalable and accurate long-read assembly via adaptive k -mer weighting and repeat separation. Genome Res 2017; 27: 722–736.

[13] Kolmogorov M, Yuan J, Lin Y, et al. Assembly of long, error-prone reads using repeat graphs. Nat Biotechnol 2019; 37: 540–546.

[14] Chen Y, Nie F, Xie S-Q, et al. Efficient assembly of nanopore reads via highly accurate and intact error correction. Nat Commun 2021; 12: 60.

[15] Li H. Minimap and miniasm: fast mapping and de novo assembly for noisy long sequences. Bioinformatics 2016; 32: 2103–2110.

[16] Shafin K, Pesout T, Lorig-Roach R, et al. Nanopore sequencing and the Shasta toolkit enable efficient de novo assembly of eleven human genomes. Nat Biotechnol 2020; 38: 1044–1053.

[17] Rao J, Peng L, Liang X, et al. Performance of copy number variants detection based on whole-genome sequencing by DNBSEQ platforms. BMC Bioinformatics 2020; 21: 518.

[18] Patterson J, Carpenter EJ, Zhu Z, et al. Impact of sequencing depth and technology on de novo RNA-Seq assembly. BMC Genomics 2019; 20: 604.

[19] Kozarewa I, Ning Z, Quail MA, et al. Amplification-free Illumina sequencing-library preparation facilitates improved mapping and assembly of (G+C)-biased genomes. Nat Methods 2009; 6: 291–5.

[20] Marx V. Method of the year: long-read sequencing. Nat Methods 2023; 20: 6–11.

[21] Dominik Stanojević, Dehui Lin, Paola Florez de Sessions, et al. Telomere-to-telomere phased genome assembly using error-corrected Simplex nanopore reads. BioRxiv.

[22] Nie F, Ni P, Huang N, et al. De novo diploid genome assembly using long noisy reads. Nat Commun 2024; 15: 2964.

[23] Holley G, Beyter D, Ingimundardottir H, et al. Ratatosk: hybrid error correction of long reads enables accurate variant calling and assembly. Genome Biol 2021; 22: 28.

[24] Lorig-Roach R, Meredith M, Monlong J, et al. Phased nanopore assembly with Shasta and modular graph phasing with GFAse. Genome Res. Epub ahead of print 16 April 2024. DOI: 10.1101/gr.278268.123.

[25] Lorig-Roach R, Meredith M, Monlong J, et al. Phased nanopore assembly with Shasta and modular graph phasing with GFAse. bioRxiv. Epub ahead of print 22 February 2023. DOI: 10.1101/2023.02.21.529152.

[26] Mikheenko A, Prjibelski A, Saveliev V, et al. Versatile genome assembly evaluation with QUAST-LG. Bioinformatics 2018; 34: i142–i150.

[27] Rhie A, Walenz BP, Koren S, et al. Merqury: reference-free quality, completeness, and phasing assessment for genome assemblies. Genome Biol 2020; 21: 245.

[28] Cheng H, Concepcion GT, Feng X, et al. Haplotype-resolved de novo assembly using phased assembly graphs with hifiasm. Nat Methods 2021; 18: 170–175.

[29] Li H. New strategies to improve minimap2 alignment accuracy. Bioinformatics 2021; 37: 4572–4574.

[30] Li H. Aligning sequence reads, clone sequences and assembly contigs with BWA-MEM.

[31] Shen W, Le S, Li Y, et al. SeqKit: A Cross-Platform and Ultrafast Toolkit for FASTA/Q File Manipulation. PLoS One 2016; 11: e0163962.

[32] Wang O, Chin R, Cheng X, et al. Efficient and unique cobarcoding of second-generation sequencing reads from long DNA molecules enabling cost-effective and accurate sequencing, haplotyping, and de novo assembly. Genome Res 2019; 29: 798–808.

[33] Höjer P, Frick T, Siga H, et al. BLR: a flexible pipeline for haplotype analysis of multiple linked-read technologies. Nucleic Acids Res 2023; 51: e114–e114.

[34] Zook JM, Catoe D, McDaniel J, et al. Extensive sequencing of seven human genomes to characterize benchmark reference materials. Sci Data 2016; 3: 160025.

[35] Sereika M, Kirkegaard RH, Karst SM, et al. Oxford Nanopore R10.4 long-read sequencing enables the generation of near-finished bacterial genomes from pure cultures and metagenomes without short-read or reference polishing. Nat Methods 2022; 19: 823–826.

[36] Yu W, Luo H, Yang J, et al. Comprehensive assessment of 11 de novo HiFi assemblers on complex eukaryotic genomes and metagenomes. Genome Res 2024; 34: 326–340.

[37] Peters BA, Kermani BG, Sparks AB, et al. Accurate whole-genome sequencing and haplotyping from 10 to 20 human cells. Nature 2012; 487: 190–195.

[38] Koren S, Walenz BP, Berlin K, et al. Canu: scalable and accurate long-read assembly via adaptive k -mer weighting and repeat separation. Genome Res 2017; 27: 722–736.

[39] Huang Y-T, Liu P-Y, Shih P-W. Homopolish: a method for the removal of systematic errors in nanopore sequencing by homologous polishing. Genome Biol 2021; 22: 95.

[40] Walker BJ, Abeel T, Shea T, et al. Pilon: An Integrated Tool for Comprehensive Microbial Variant Detection and Genome Assembly Improvement. PLoS One 2014; 9: e112963.

[41] Bankevich A, Bzikadze A V., Kolmogorov M, et al. Multiplex de Bruijn graphs enable genome assembly from long, high-fidelity reads. Nat Biotechnol 2022; 40: 1075–1081.

