## Supplementary Figures for "Pre-Assembly NGS Correction of ONT Reads Achieves HiFi-Level Assembly Quality"

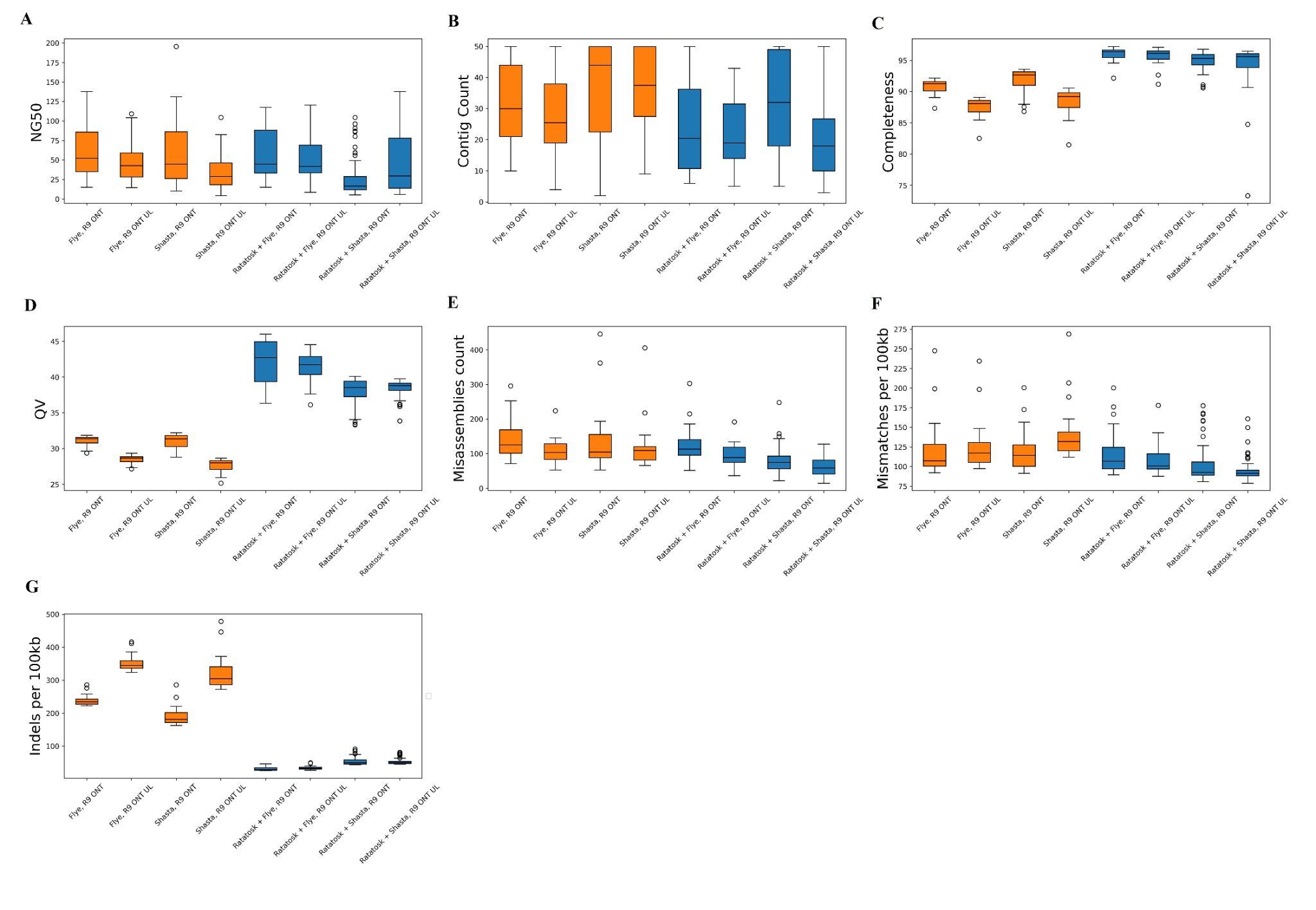

**Figure 1. Quality statistics distribution of assemblies on chromosomes 1 to 22 of HG002**. Solid lines for corrected ONT-based assemblies, dashed for not corrected ONT-based assemblies. **A.** NG50, **B.** Contig Count, **C.** K-mer based completeness, **D.** QV score. **E.** Overall count of misassembles **F.** Mismatches per 100kb **G.** Indels per 100kb.

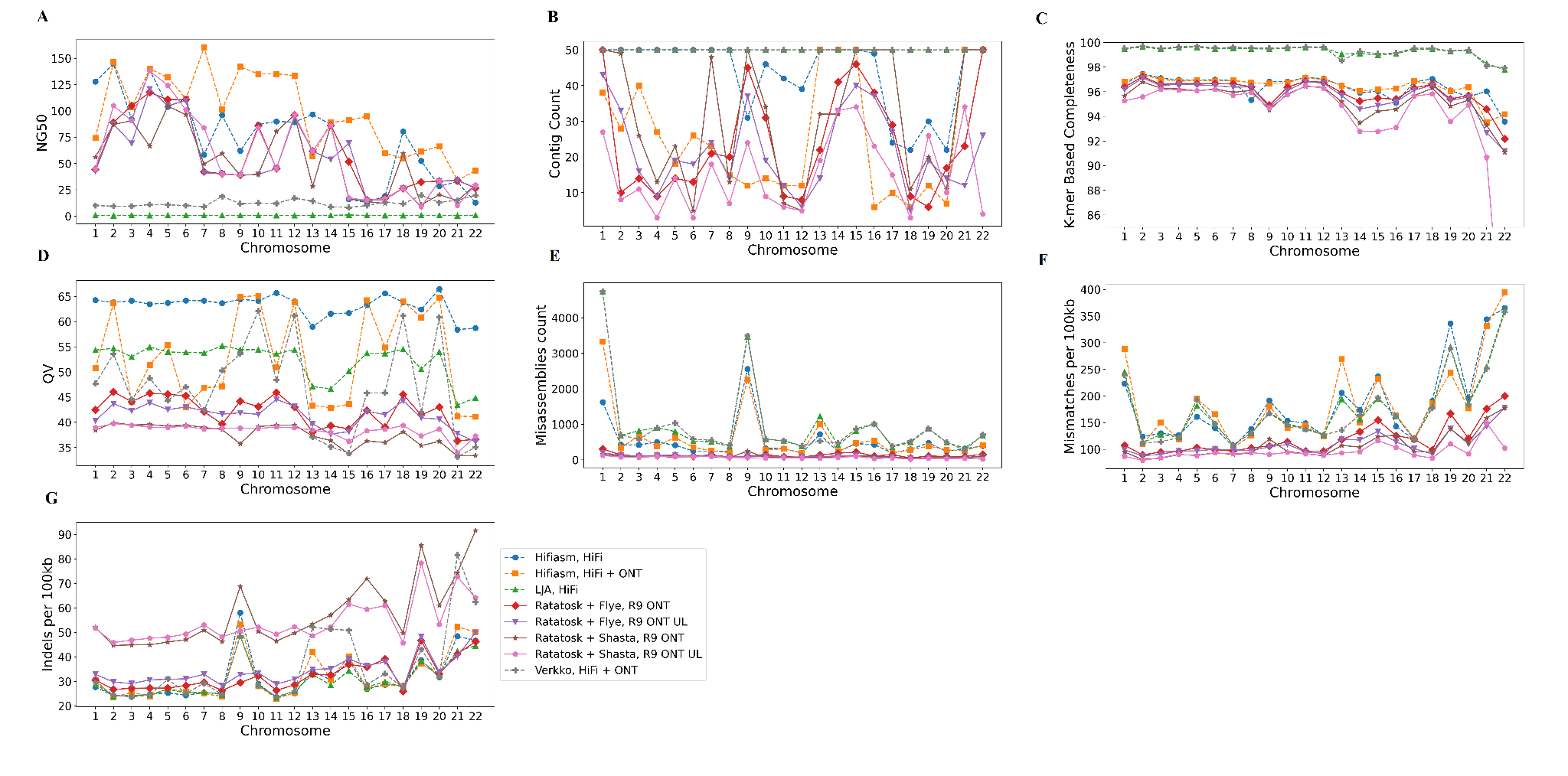
**Figure 2. Quality statistics distribution of assemblies on chromosomes 1 to 22 of HG002**. Solid lines for ONT + NGS based assemblies, dashed for HiFi or HiFi + ONT based assemblies. **A.** NG50, **B.** Contig Count, **C.** K-mer based completeness, **D.** QV score. **E.** Overall count of misassembles **F.** Mismatches per 100kb **G.** Indels per 100kb.

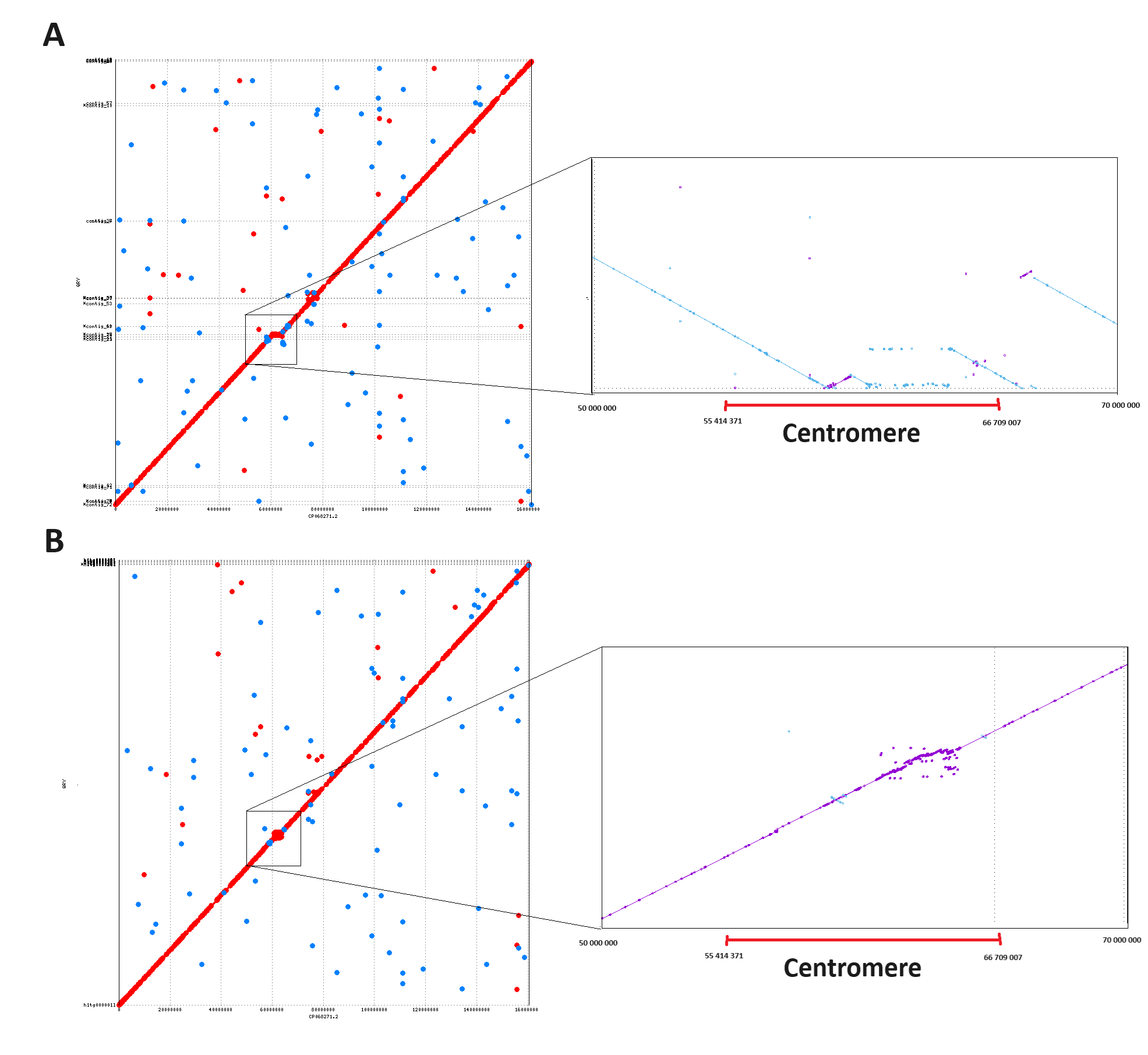

**Figure 3. Mummerplot of assemblies of chr7 vs CHM13 reference**. **A.** Ratatosk + Flye with NGS + R9 ONT assembly. **B.** Hifiasm with HiFi + R9 ONT UL assembly.

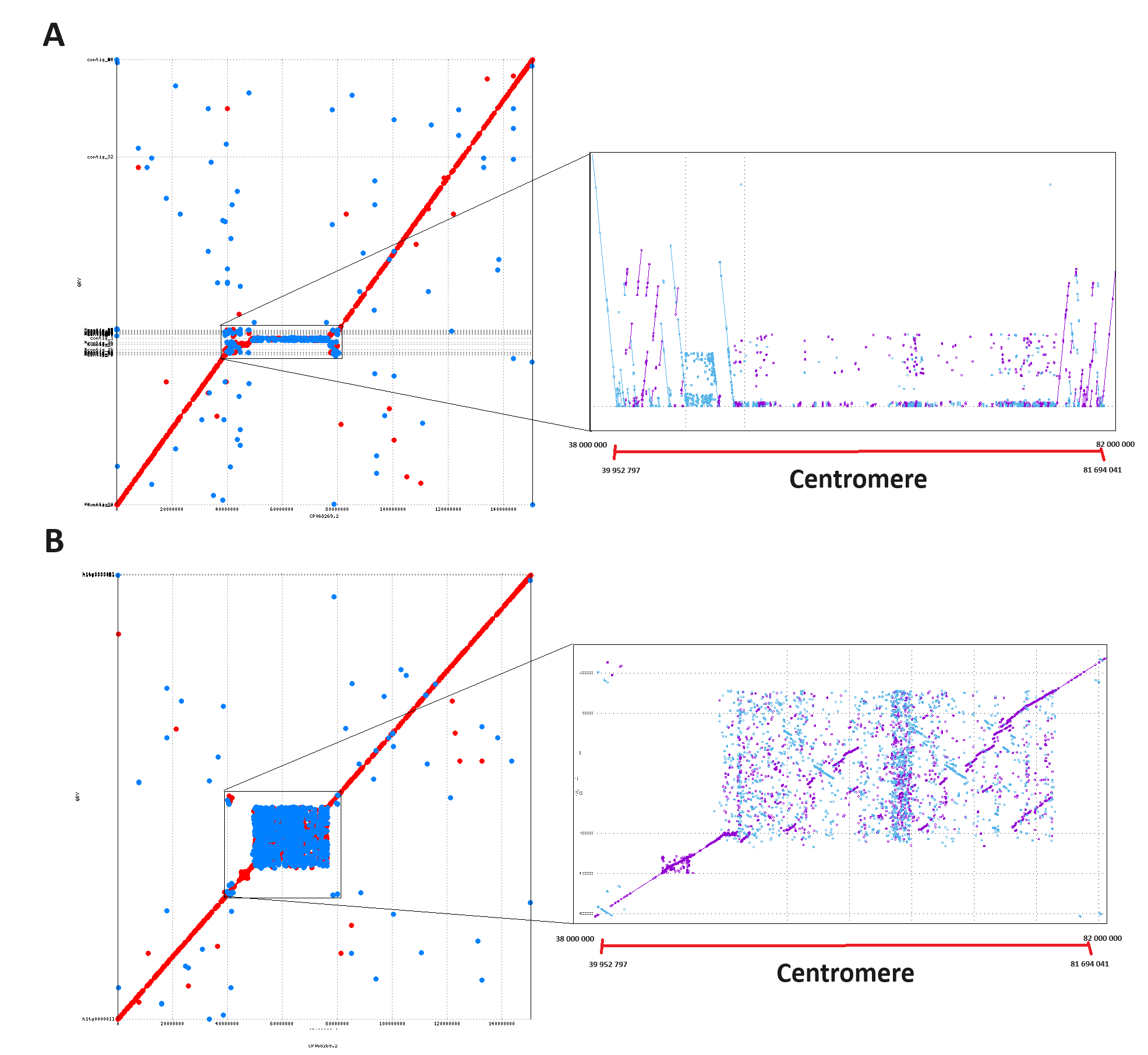

**Figure 4. Mummerplot of assemblies of chr9 vs CHM13 reference**. **A.** Ratatosk + Flye with NGS + R9 ONT assembly. **B.** Hifiasm with HiFi + R9 ONT UL assembly.

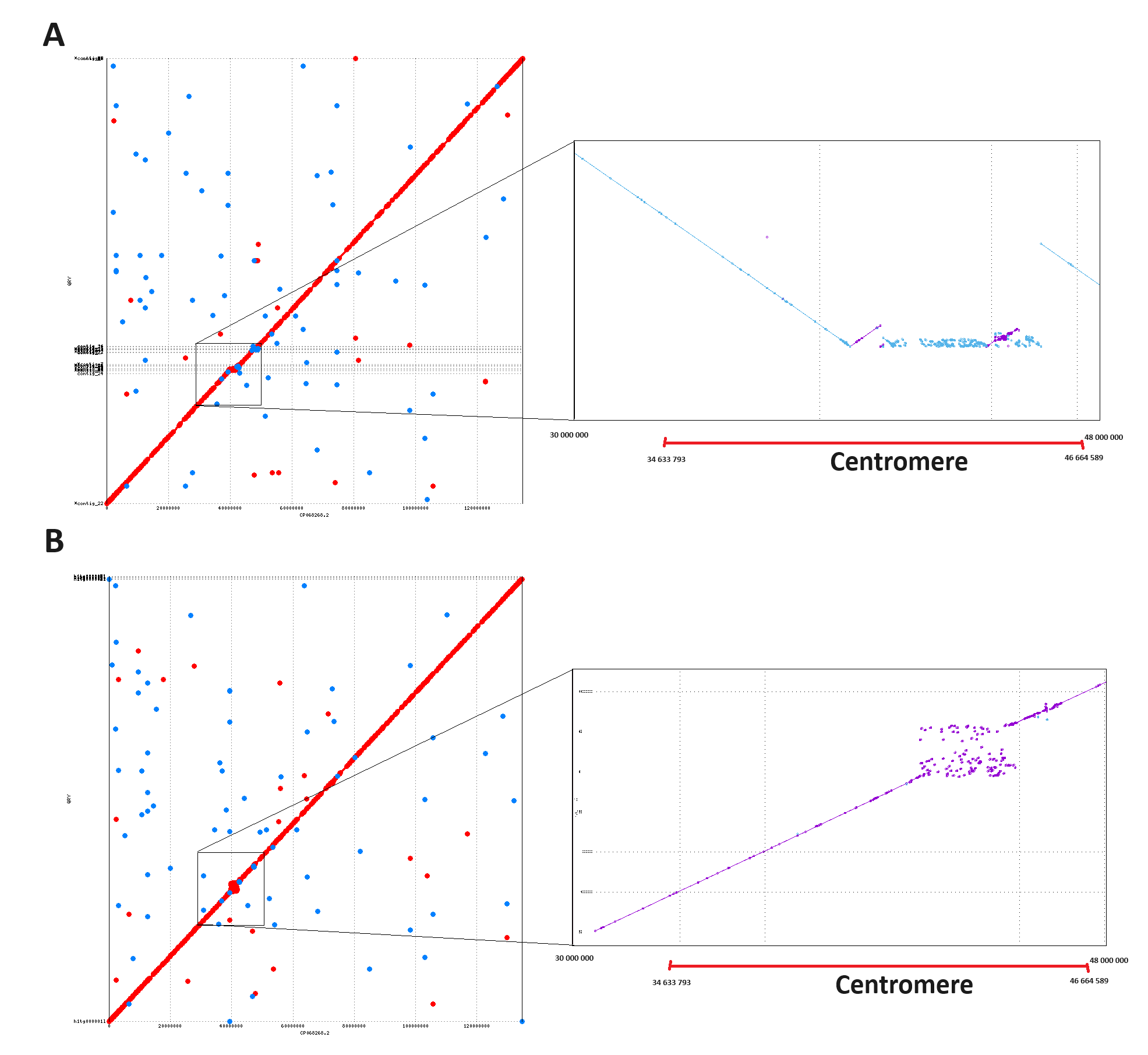

**Figure 5. Mummerplot of assemblies of chr10 vs CHM13 reference**. **A.** Ratatosk + Flye with NGS + R9 ONT assembly. **B.** Hifiasm with HiFi + R9 ONT UL assembly.

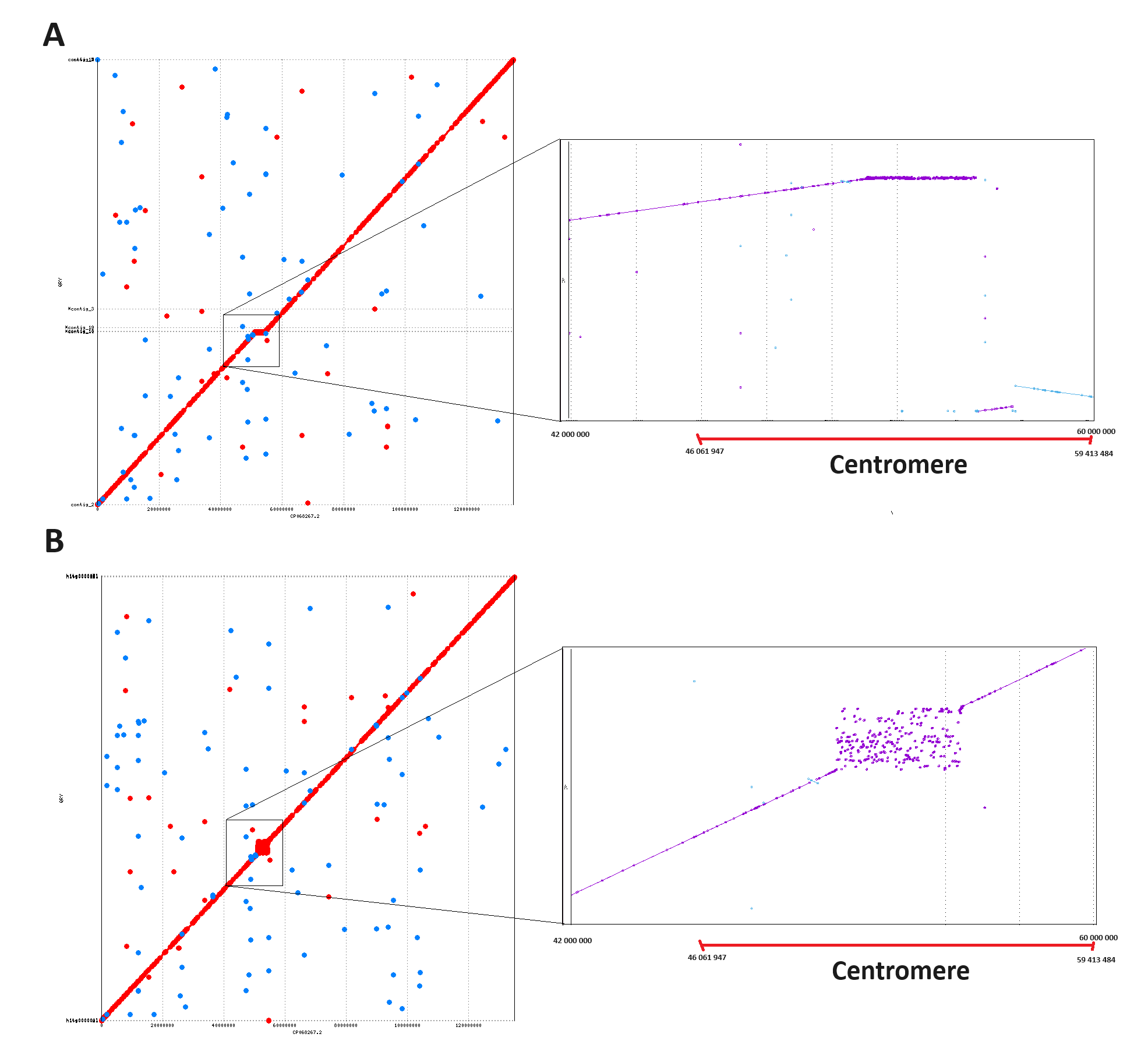

**Figure 6. Mummerplot of assemblies of chr11 vs CHM13 reference**. **A.** Ratatosk + Flye with NGS + R9 ONT assembly. **B.** Hifiasm with HiFi + R9 ONT UL assembly.

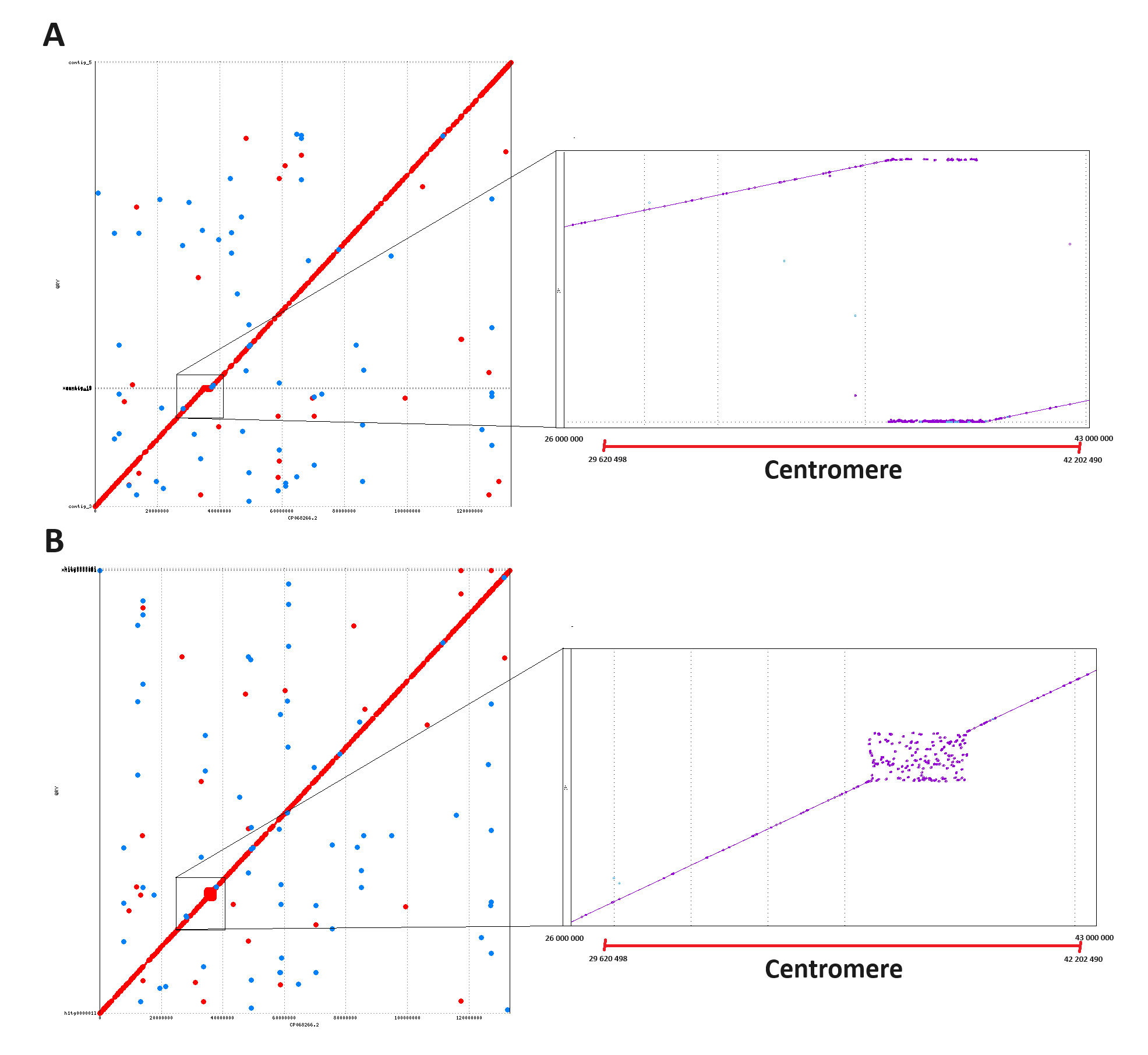

**Figure 7. Mummerplot of assemblies of chr12 vs CHM13 reference**. **A.** Ratatosk + Flye with NGS + R9 ONT assembly. **B.** Hifiasm with HiFi + R9 ONT UL assembly.

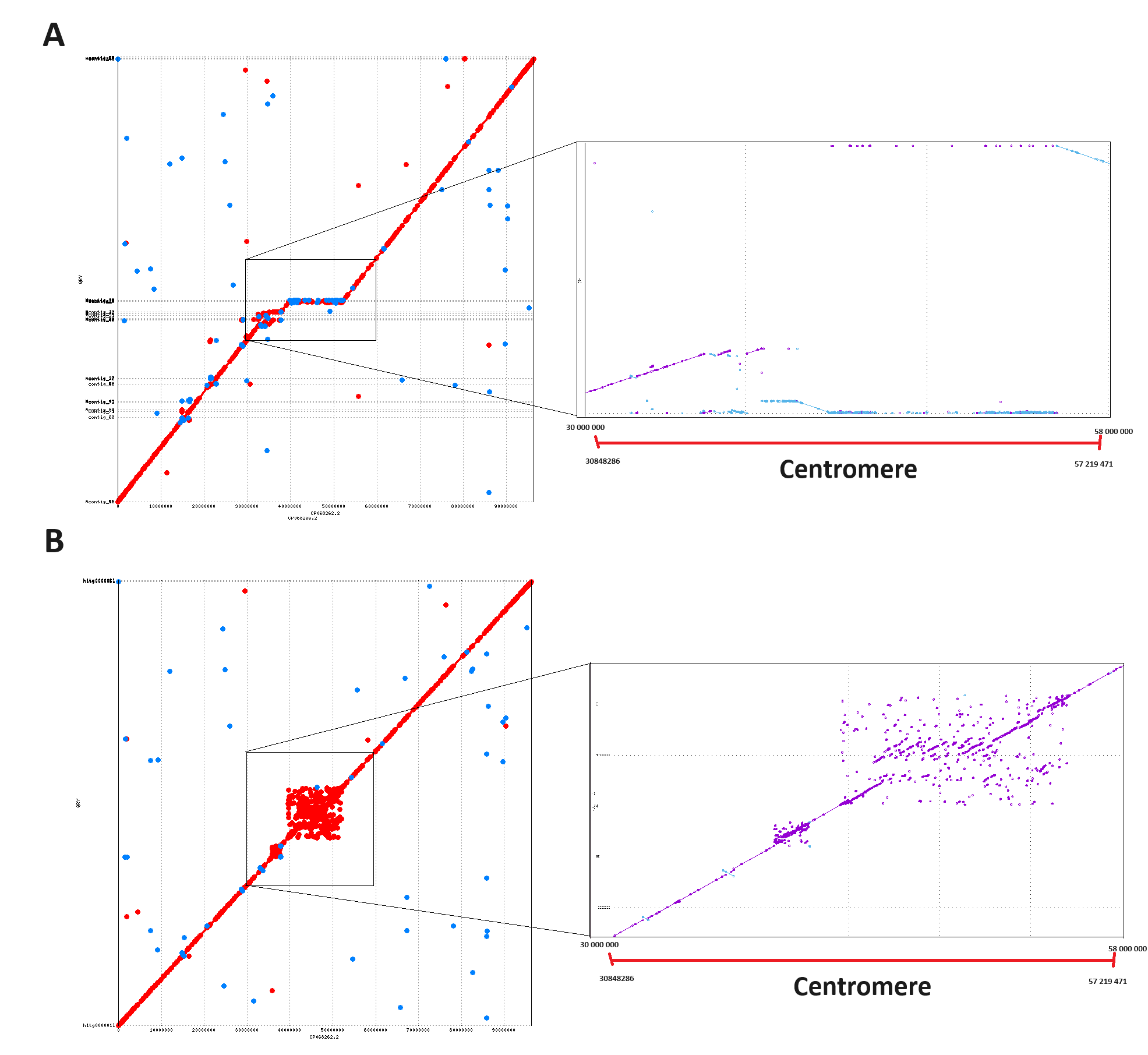
**Figure 8. Mummerplot of assemblies of chr16 vs CHM13 reference**. **A.** Ratatosk + Flye with NGS + R9 ONT assembly. **B.** Hifiasm with HiFi + R9 ONT UL assembly.

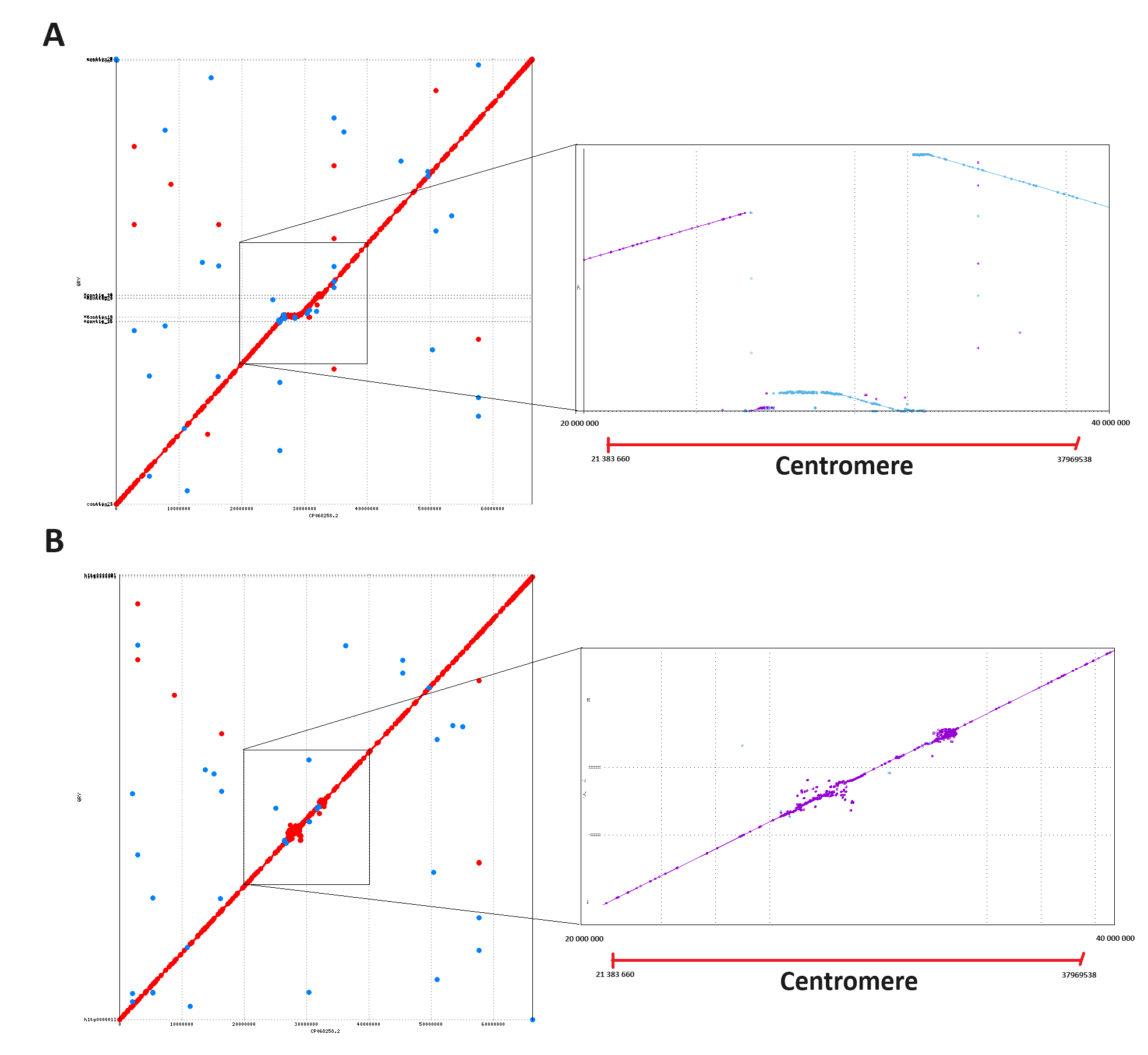

**Figure 9. Mummerplot of assemblies of chr20 vs CHM13 reference**. **A.** Ratatosk + Flye with NGS + R9 ONT assembly. **B.** Hifiasm with HiFi + R9 ONT UL assembly.

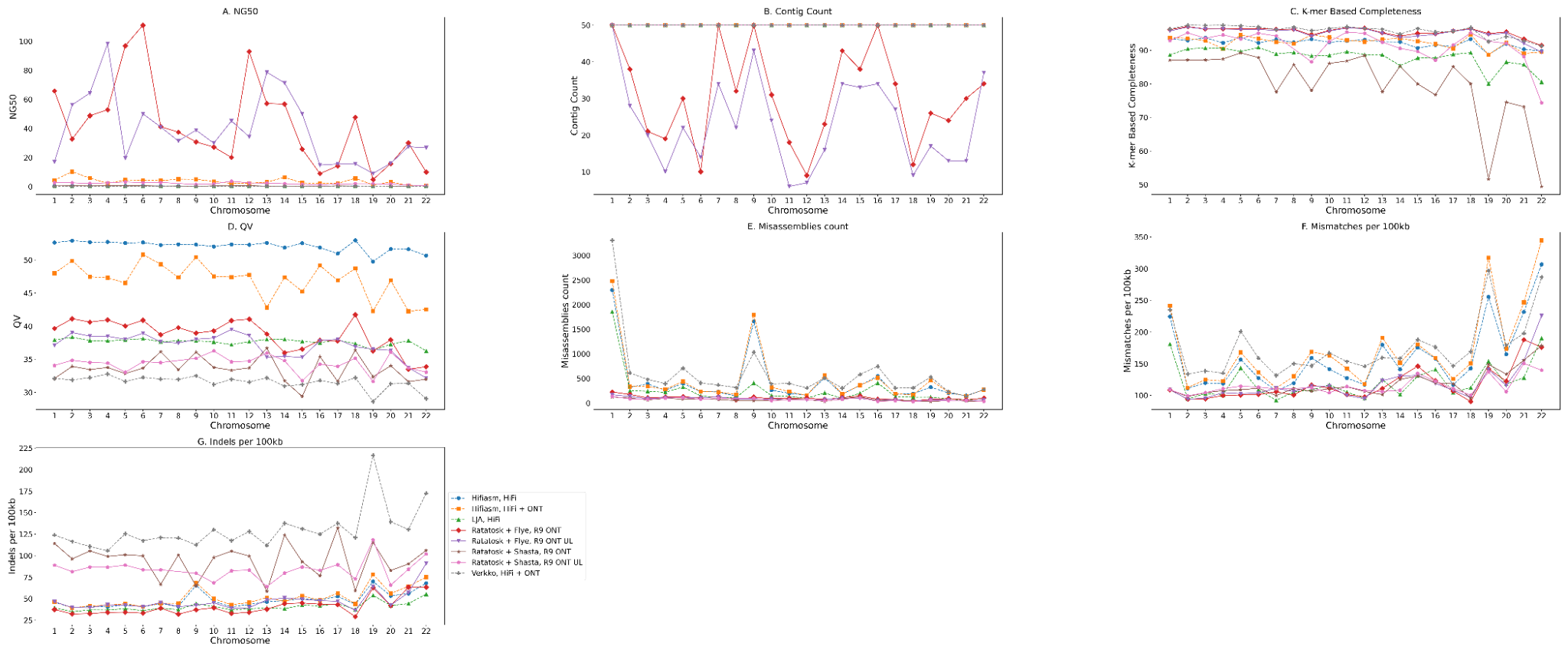

**Figure 10. Average values of quality statistics of low coverage assemblies on chromosomes 1 to 22 of HG002**. Solid lines for ONT + NGS based assemblies, dashed for HiFi or HiFi + ONT based assemblies. **A.** NG50, **B.** Contig Count, **C.** K-mer based completeness, **D.** QV score. **E.** Overall count of misassembles **F.** Mismatches per 100kb **G.** Indels per 100kb.

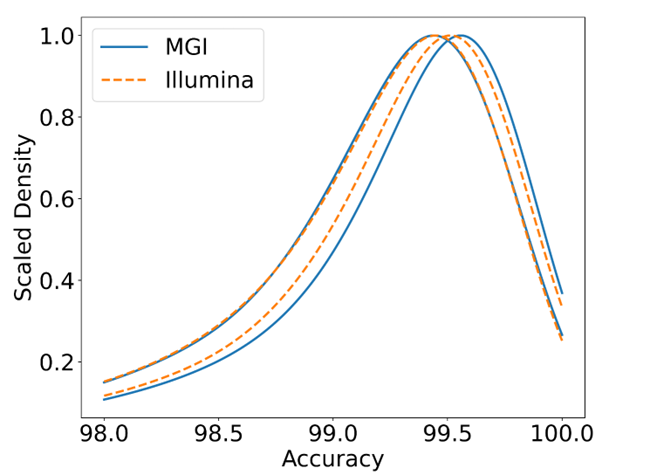

**Figure 11. Empirical accuracy of Ratatosk corrected ONT reads with NGS**. Scaled density of empirical accuracy of corrected ONT reads by NGS short reads. Blue lines - MGI, orange lines - Illumina NGS short reads. Only chromosome 20 was utilized for this evaluation.
